# Exercise-induced sweat suppresses depression-like behaviors in mice

**DOI:** 10.64898/2026.06.02.728455

**Authors:** Huilin Yao, Sisi Xing, Maodi Liang, Qintong Fei, Jing Cao, Tiantian Liang, Bojun Yang, Chunmei Cui, Hua Liu, Qinghua Cui

## Abstract

Depression represents a major global public health challenge. Current antidepressant therapies are often limited by significant side effects, delayed onset of action, and suboptimal efficacy, creating an urgent need for the development of novel interventions. In this study, using a transcriptome-based bioinformatics pipeline, we found that the gene signature of depression is significantly anti-correlated with that of exercise-induced sweat (EIS), suggesting that EIS may have the potential to treat depression. To validate this prediction, we first established a lipopolysaccharide-induced mouse model of depression and then evaluated the therapeutic effects of EIS in comparison with the positive control fluoxetine. Our results show that EIS treatment significantly suppresses depressive and anxiety-like behaviors, as evidenced by increased sucrose preference in the sucrose preference test, reduced immobility in the tail suspension test, and enhanced exploratory activity in the elevated plus maze. Notably, high-dose EIS intervention achieved therapeutic efficacy comparable to, or even better than, that of fluoxetine. This study identifies EIS as a novel bioactive intervention and an innovative strategy for treating depression, and opens a new avenue in sports medicine by investigating the potential beneficial effects of EIS.

## Introduction

Depression represents a major global public health challenge, characterized by persistent low mood, anhedonia, and cognitive impairment, which are associated with high rates of disability and mortality^1^. Although selective serotonin reuptake inhibitors (SSRIs) such as fluoxetine are widely prescribed for major depressive disorder, their clinical use is limited by delayed therapeutic onset, incomplete remission, and adverse side effects^2^. In addition, access to treatment for many people with depression is limited, with only 51% treatment coverage for high income countries and 20% for low and lower-middle income countries^3^. Therefore, there is an urgent need to identify new, effective and safe treatment strategies.

Increasing evidence highlights the critical role of neuroinflammation in the pathogenesis of depression. Both clinical and experimental studies show that depression is associated with elevated pro-inflammatory cytokines and the activation of immune-related signaling pathways^4-6^. Lipopolysaccharide (LPS), a well-established systemic inflammatory inducer, is commonly used to model neuroinflammation-related depression in animals, resulting in behavioral changes such as anhedonia and despair^7,8^. These findings underscore neuroinflammation as a key therapeutic target for depression^9^.

Lifestyle factors, particularly physical exercise, are known to have positive effects on mental health, reducing depressive symptoms and improving emotional regulation^10-13^. Traditionally, these benefits are attributed to central mechanisms, including enhanced neurogenesis, neurotransmitter modulation, and anti-inflammatory signaling^14,15^. However, exercise also triggers systemic physiological changes, including the release of various bioactive substances into body fluids, which may contribute to its overall therapeutic effects^16^.

Sweat is an easily accessible biological fluid. Emerging evidence indicates that sweat is not merely a metabolic waste product but a complex system containing diverse bioactive molecules, such as cytokines, peptides, and immune-related factors^17-20^. These components have been reported to possess antimicrobial, anti-inflammatory, and tissue-regulatory functions, suggesting that sweat may have biological roles beyond its traditional functions. We previously revealed that exercise-induced sweat (EIS) promotes wound healing in diabetic foot ulcers^21^. However, it remains largely unknown whether EIS can directly mediate the beneficial effects of exercise in the context of neuroinflammation and depression.

Transcriptomics provides a powerful strategy for evaluating the therapeutic potential of complex biological systems. By comparing gene expression signatures induced by specific interventions with those associated with disease, one can predict potential therapeutic effects without the need to isolate individual components^22,23^. This approach is particularly suitable for studying heterogeneous biological fluids like sweat, as it captures their comprehensive biological effects.

In this study, using the above transcriptome-based strategy, we found that the gene signature of depression is significantly anti-correlated with that of sweat induced by moderate-intensity aerobic exercise, suggesting that EIS may defense depression. Then we validated the above prediction and hypothesis using mouse experiments. The results showed that EIS treatment significantly increased sucrose preference in the sucrose preference test (SPT) and reduced immobility time in the tail suspension test (TST), indicating improvements in anhedonia and behavioral despair. Additionally, anxiety-like behaviors were alleviated, as evidenced by increased time spent in the center region and open arms of the elevated plus maze (EPM). Our study suggests that as a natural product, EIS could inhibit the progression of depression, providing a safer, more cost-effective, and more efficient option for the clinical management of depression.

## Materials and Methods

### Participants in moderate-intensity aerobic exercise

As described in our previous study, this study enrolled five healthy adult female volunteers, aged 22–30 years, with a body mass index (BMI) of 18–25 kg/m² and no history of systemic diseases^21,24^. In the month prior to enrollment, no medications known to affect sweating or inflammatory responses were taken by any participants. Following a standardized protocol, all participants underwent a thorough pre-examination screening. This included the collection of basic demographic information (Supplementary File S1), completion of a Physical Activity Readiness Questionnaire (PAR-Q), and baseline medical checks such as blood pressure measurements and routine blood tests.^25^. This screening ensured that every participant could safely complete the moderate-intensity aerobic exercise session for sweat collection.

The study purpose, procedures, sample collection methods, potential risks and benefits, and privacy safeguards were explained to all participants in clear language. Written informed consent was obtained from each participant. Throughout the study, biological sample collection and data handling were conducted in strict accordance with ethical requirements. Participants were informed that they could withdraw at any time without any negative consequences.All study procedures were approved by the Ethics Committee of Wuhan Sports University (Approval No.L2025089) prior to study initiation.

### Exercise protocol

As none of the participants had a history of regular exercise training, each individual’s maximum heart rate (HRmax) was estimated using the Tanaka formula:: HRmax = 208 − (0.7 × age)^26^. Based on this, the target exercise intensity was set at 60%–80% of the individual’s HR_max to ensure a moderate-intensity range. Based on participants’ ages, the resulting target heart rate range was approximately 116–138 beats per minute.

To minimize the influence of potential confounding factors on sweat secretion and metabolic state, all experimental sessions were scheduled outside the participants’ menstrual periods. For 24 hours prior to the exercise intervention, a standardized pre-experimental protocol was strictly followed. This included abstaining from alcohol, coffee, strong tea, and other caffeinated beverages, as well as avoiding high-sugar, high-fat, spicy, or stimulant-rich foods. Additionally, a minimum 2-hour fast was required before the experiment. These measures **were** implemented to reduce metabolic variability caused by dietary differences and to prevent interference with subsequent biofluid-related experiments.The specific exercise protocol was as follows: Participants first performed a 5-minute progressive warm-up on a treadmill, followed by 30 minutes of continuous running while maintaining their individual target heart rate range^27^. Throughout the entire session, heart rate was monitored continuously and in real time using a Polar H10 heart rate sensor to ensure the exercise intensity consistently met the preset requirements.

### Sweat collection and sample processing

Sweat was collected using an absorbent filter paper patch method. Prior to sampling, the skin on the participant’s back was cleansed with alcohol and deionized water, then dried with sterile gauze. Two pieces of absorbent filter paper (Whatman™ quantitative filter paper, Grade 1825-025, Cytiva, UK), each 25 mm in diameter, were placed on the skin. These were covered with a waterproof transparent medical adhesive dressing (3M™ Tegaderm™ transparent film dressing, model 1624W, 3M Health Care, USA) to secure them in place. Throughout the procedure, the researchers wore disposable gloves and handled the filter paper only with sterile forceps to minimize exogenous contamination^28^.Sweat collection lasted approximately 25Lminutes to obtain sufficient volume for subsequent analysis. During this period, the patch was visually inspected at regular intervals; if the filter paper appeared close to saturation, it was removed early to prevent sweat overflow or sample loss.

After collection, the filter paper was immediately transferred to preLcooled, nucleaseLfree centrifuge tubes. For elution, 2 mL of normal saline was added to each tube. The tubes were agitated on an orbital shaker at 4 °C and 200 rpm for 2 hours. The resulting eluate was then centrifuged at 4 °C and 4000 rpm for 5 min. The supernatant was collected and passed through a 0.22 μm syringe filter to obtain a clear sweat sample. All samples were stored in an ultra-low-temperature freezer (−80 °C) until further analysis.During storage, water evaporation may lead to overestimation of solute concentration in sweat samples^29^. To reduce this bias, following previous recommendations^29,30^L, samples were sealed with impermeable Parafilm®LM film to improve sample stability and the reliability of analytical results.

### Cell culture and treatment

In order to obtain the transcriptome induced by EIS, we select the A549 cell line as the tool cell and treated A549 with EIS and the control. The A549 cell line was routinely cultured in high-glucose DMEM supplemented with 10% fetal bovine serum and 1% penicillin-streptomycin, and maintained at 37 °C in a humidified atmosphere with 5% COL. Cells in the logarithmic growth phase were seeded into 6-well plates. Twelve hours after seeding in 6-well plates, the medium was replaced with fresh medium containing either EIS (HE) or an equal volume of saline control (Hcon). The cells were then incubated for 48 hours. Three independent biological replicates were performed for each experimental group.

### RNA sequencing and transcriptomic Analysis

Total RNA was isolated from A549 cells using Trizol reagent (Invitrogen). RNA quality and integrity were assessed with an Agilent 2100 BioAnalyzer and a Qubit Fluorometer; only samples with an RNA integrity number (RIN) > 7.0 and a 28S:18S ratio > 1.8 were used for downstream processing.

RNA-seq libraries were constructed by CapitalBio Technology (Beijing, China) using the NEB Next Ultra RNA Library Prep Kit for Illumina. Briefly, poly(A)+ mRNA was enriched from 1 µg of total RNA, fragmented, and converted into double-stranded cDNA. After end repair, A-tailing, and adapter ligation, libraries were amplified by PCR. Final library quality and concentration were validated using an Agilent 2100 Bioanalyzer and the KAPA Library Quantification kit. Libraries were then subjected to paired-end 150 bp sequencing on an Illumina NovaSeq platform.

For data analysis, raw read quality was assessed with FastQC (v0.11.5) and adapters/low-quality bases were trimmed using NGSQC (v2.3.3). Cleaned reads were aligned to the human reference genome (hg38) using HISAT2 (v2.1.0). Transcript assembly and gene-level expression quantification were performed with StringTie (v1.3.3b). Using the same pipeline as our previous studies, the genes with a fold change (FC) ≥ 1.50 or ≤ 0.67 were determined as the gene signature induced by EIS. Next, an in-house computer program was used to predict the potential protective roles of EIS based on the reverse transcriptomics strategy.

### Experimental Animals and Ethics

Male C57BL/6J mice (8 weeks old) were used in this study and were purchased from the Hubei Provincial Center for Disease Control and Prevention (Hubei, China). Given the clear sex differences in neuroinflammation and depression-like behaviors, only male mice were included to avoid potential confounding effects of hormonal fluctuations in females^31,32^. All animals were housed under controlled environmental conditions (temperature: 22-24 °C; relative humidity: 55 ± 3%; 12 h light/dark cycle, lights on from 08:00 to 20:00) with ad libitum access to standard chow and autoclaved water. All experimental procedures were conducted in accordance with the guidelines of the Ethics Committee of Wuhan Sports University (ECWSU, Wuhan, China; approval ID: S0087-20250908-01). Every effort was made to minimize animal suffering and to reduce the number of animals used while ensuring the reliability of the experimental results. At the end of the experiment, mice were euthanized. Whole brain tissues for histopathological analysis and other tissues for biochemical assays were rapidly collected, snap-frozen in liquid nitrogen, and stored for subsequent analyses.

### Experimental Design and Intervention

To establish the neuroinflammation-induced depression model, mice were subjected to daily intraperitoneal (i.p.) injections of LPS (1 mg/kg, 0.2 ml) for five consecutive days. These LPS-treated mice were randomly assigned to four experimental groups: a vehicle control group (Vehicle), a low-dose sweat group (Sweat-low), a high-dose sweat group (Sweat-high), and a fluoxetine positive control group. For therapeutic interventions, mice in the Vehicle, Sweat-low, and Sweat-high groups received oral gavage of normal saline, 1:1 diluted sweat, or undiluted sweat (0.2 ml per dose), respectively. This gavage was performed four times daily at 2-hour intervals, commencing at 10:00 a.m. Alternatively, the fluoxetine group received an oral gavage of fluoxetine hydrochloride (10 mg/kg). Across all groups, the daily LPS injection was administered 1 h following the final gavage treatment.

### Sucrose Preference Test (SPT)

The sucrose preference test was performed to assess anhedonia-like behavior. After the final intraperitoneal injection, mice were deprived of food for 6 h and individually housed. Subsequently, mice were habituated to two identical bottles containing either 1% (w/v) sucrose solution or tap water for 48 h. The positions of the bottles were switched every 24 h to prevent side bias. During the test phase, mice were given free access to both solutions for 12 h. Bottle positions were randomized at the start of the test and switched after 6 h^33^. Fluid consumption was measured, and sucrose preference was calculated as follows: sucrose preference (%) = (sucrose intake / total fluid intake) × 100.

### Elevated Plus Maze (EPM)

The elevated plus maze test was used to evaluate anxiety-like behavior. The apparatus consisted of two open arms (50 × 10 cm) and two closed arms of the same size with 15-cm-high walls, connected by a central platform (10 × 10 cm), and elevated 50 cm above the floor. Each mouse was placed in the center of the maze facing an open arm and allowed to explore freely for 5 min. The time spent in the open arms and the number of entries into each arm were recorded and analyzed. The testing environment was maintained under low, uniform illumination, and the experimenter remained out of the animal’s sight during testing.

### Tail Suspension Test (TST)

The tail suspension test was conducted to assess behavioral despair. Mice were suspended by the tail using adhesive tape placed approximately 1–1.5 cm from the tip of the tail. The tape was attached to a horizontal bar, with the head of the mouse positioned at least 20 cm above the surface. Each session lasted for 6 min, and the immobility time during the last 4 min was recorded for analysis^34^.

### Open Field Test (OFT)

Mice were placed in the center of an open-field arena (44 cm × 44 cm × 44 cm) and allowed to explore freely for 5 min^35^. Locomotor activity was recorded using a camera positioned above the apparatus.

## Results

### Overview of the study design and experimental workflow

Figure 1 shows the framework of this study. Firstly, sweat induced by moderate-intensity aerobic exercise was collected from five young healthy adult female participants using an absorbent filter paper patch method. Then the A549 cell was taken as the tool cell to be treated with the sweat (HE) and the control (saline, Hcon) for 48 hours. Next, total RNAs of A549 cells were sequenced using the next-generation RNA-sequencing technique. And then bioinformatics analysis identified the gene signature of the sweat and predicted depression as the significantly potential beneficial effects of the sweat. Finally, mouse experiments validated the prediction, that is, sweat indeed suppresses depression.

**Figure 1.**
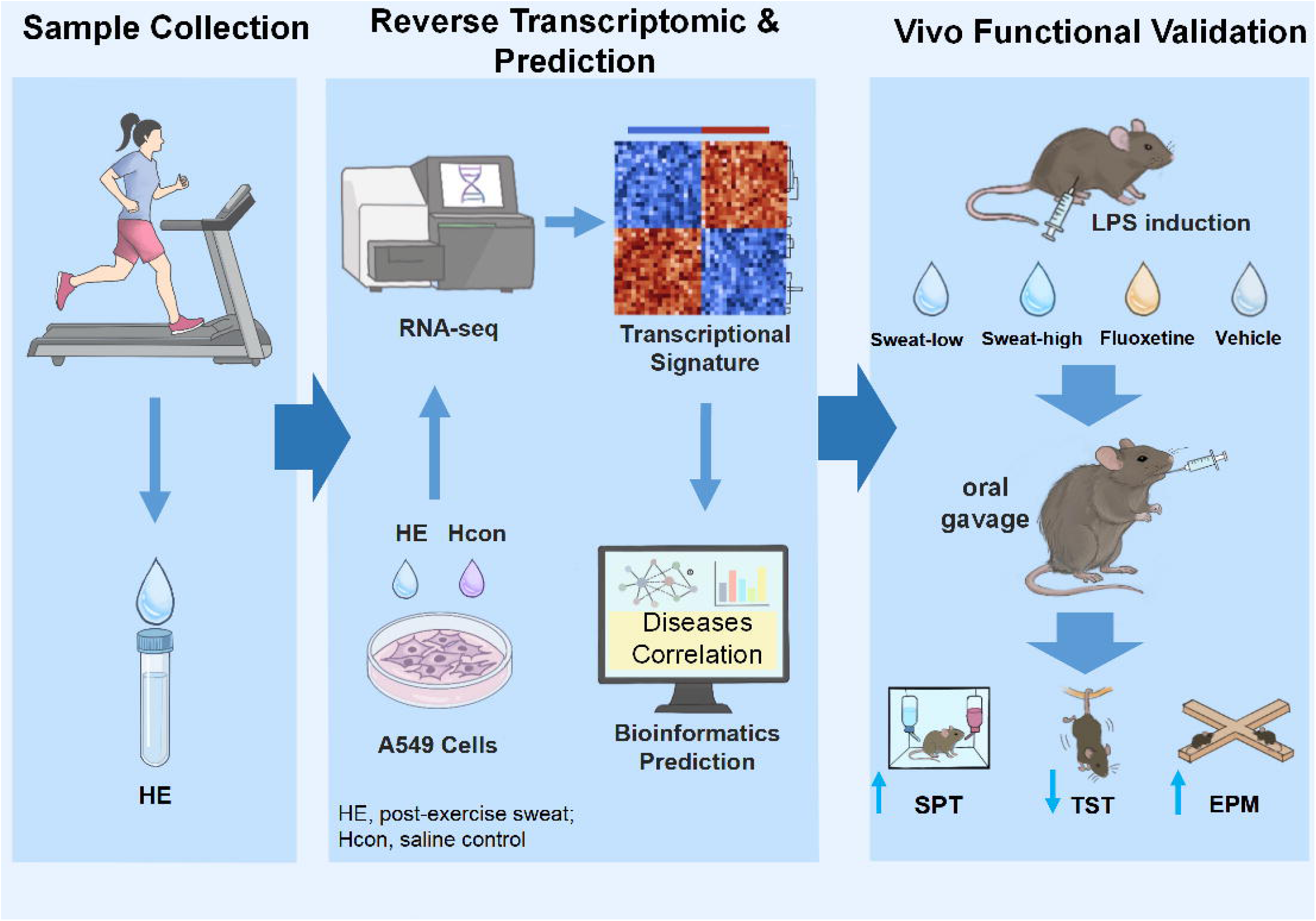
Overview of the study design and experimental workflow.

### Predicting exercise-induced sweat to treat depression

To predict whether exercise-induced sweat (EIS) may treat depression, we obtained the normalized gene expression profiles (GSE24095) of depression and control from Duric et al.’s study, which collected the postmortem hippocampus samples of 21 patients with major depressive disorder vs. 18 normal controls^36^. In their study, gene expression profiles of 15 paired of CA1 and 15 paired of dentate gyrus (DG) subregions of the hippocampus were assessed by the Microarrays Inc. Human Genomic 49K MI Ready Array platform (GPL10907). Based on the gene signatures of depression, the same cutoff (FC≥1.5) as that of EIS, the results showed that the gene signature of EIS is significantly anti-correlated with that of depression’s CA1 (OR = 0.24, p-value = 7.4e-4, Fisher’s exact test, **Figure 2A**). For DG of depression, the result is not statistically significant but has a clear tendency (OR = 0.64, p-value = 0.21, **Figure 2B**). The gene signature overlaps are shown in Supplementary File S2 & S3.

**Figure 2.**
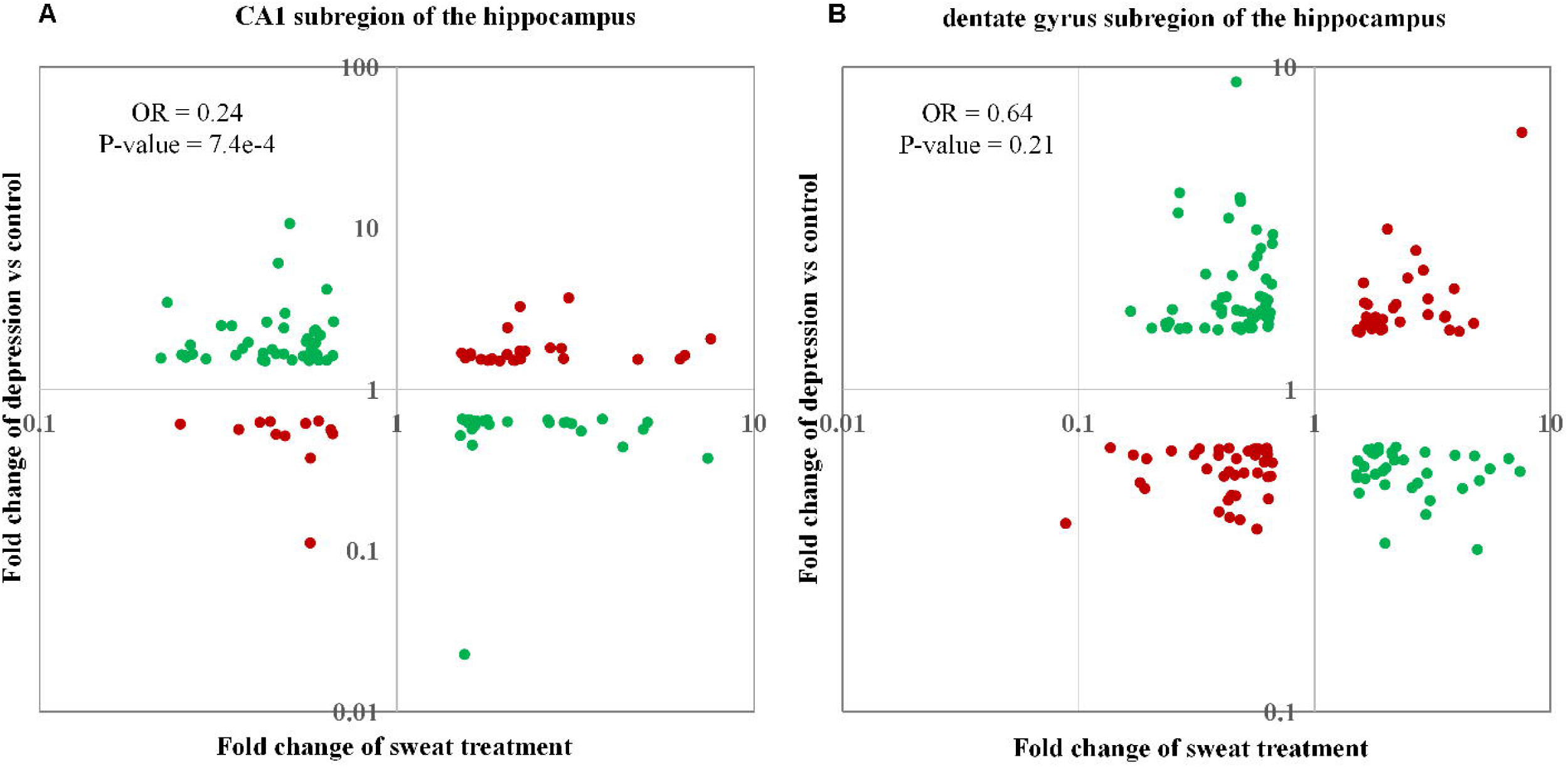
List of overlapped gene signatures of sweat treatment and depression. (a) CA1 subregion; (b) dentate gyrus subregion

### Exercise-induced sweat alleviates depressive symptoms

To investigate whether exercise-induced sweat alleviates depression-like behaviors induced by neuroinflammation, we established a neuroinflammation-induced depression model via LPS injection and administered different doses of sweat, utilizing fluoxetine as a positive control **(Figure 3A)**. Our results showed that sweat-treated mice exhibited a significant increase in sucrose preference, with an effect comparable to fluoxetine **(Figure 3B)**, indicating that sweat treatment improves LPS-induced anhedonia. Meanwhile, mice receiving sweat exhibited increased time spent in the center region **(Figure 3C, D)** and in the open arms **(Figure 3E)**, as well as the greater distances traveled in the open arms **(Figure 3F)** in the elevated plus maze test. In addition, sweat treatment significantly reduced immobility time in the tail suspension test compared with the vehicle group (**Figure 3G**), further supporting that sweat treatment improves LPS-induced behavioral despair. However, in the open field test, neither the low- nor high-dose sweat treatment significantly affected locomotor activity (**Figure 4A-B**), center time (**Figure 4C**), or the proportion of distance traveled in the center region (**Figure 4D**), although a trend toward increased center exploration was observed. Collectively, these data demonstrate that exercise-induced sweat attenuates neuroinflammation-associated behavioral deficits, alleviating depressive- and anxiety-like symptoms in mice.

**Figure 3.**
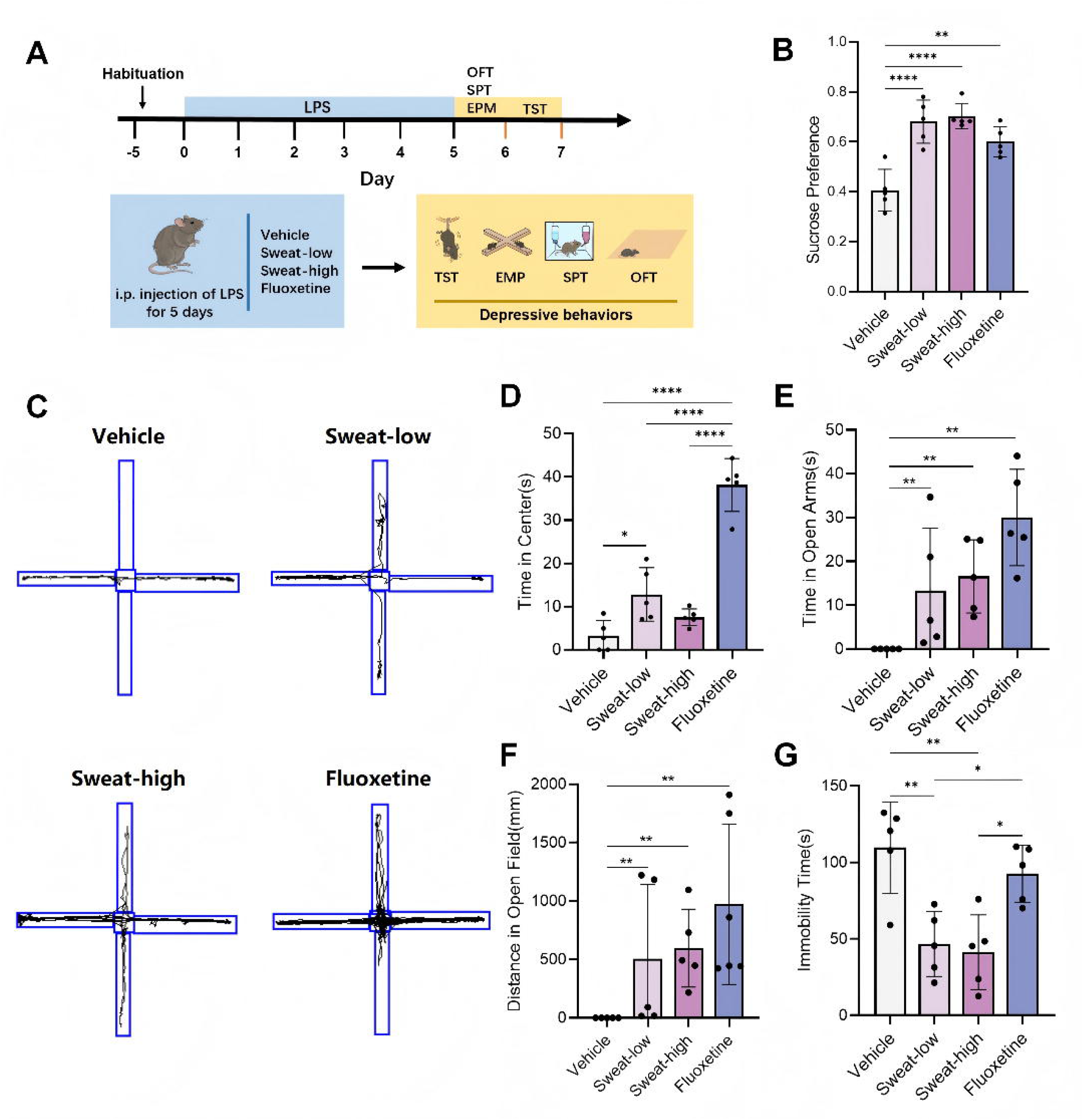
Exercise-induced sweat alleviates depressive-like and anxiety-like behaviors in LPS-induced mice. (A) Schematic representation of the experimental workflow. Mice were subjected to LPS-induced neuroinflammation and concurrently treated with vehicle, different doses of sweat, or fluoxetine. Behavioral assessments were conducted from days 5 to 7. (B) Percentage of sucrose preference in the sucrose preference test (SPT). (C–F) Elevated plus maze (EPM) test. (C) Representative locomotion tracking maps of mice in the EPM. Quantification of (D) time spent in the central platform, (E) time spent in the open arms, and (F) total distance traveled in the open arms. (G) Immobility time in the tail suspension test (TST). Data are presented as mean ± SD (n = 5 mice/group). Statistical significance was determined by one-way ANOVA followed by Tukey’s post hoc test, except for the EPM behavioral data (D-F), which were analyzed using the Kruskal-Wallis test followed by the Mann-Whitney U test due to data distribution characteristics. \**P* < 0.05, \*\**P* < 0.01, \*\*\**P* < 0.001, \*\*\*\**P* < 0.0001.

**Figure 4.**
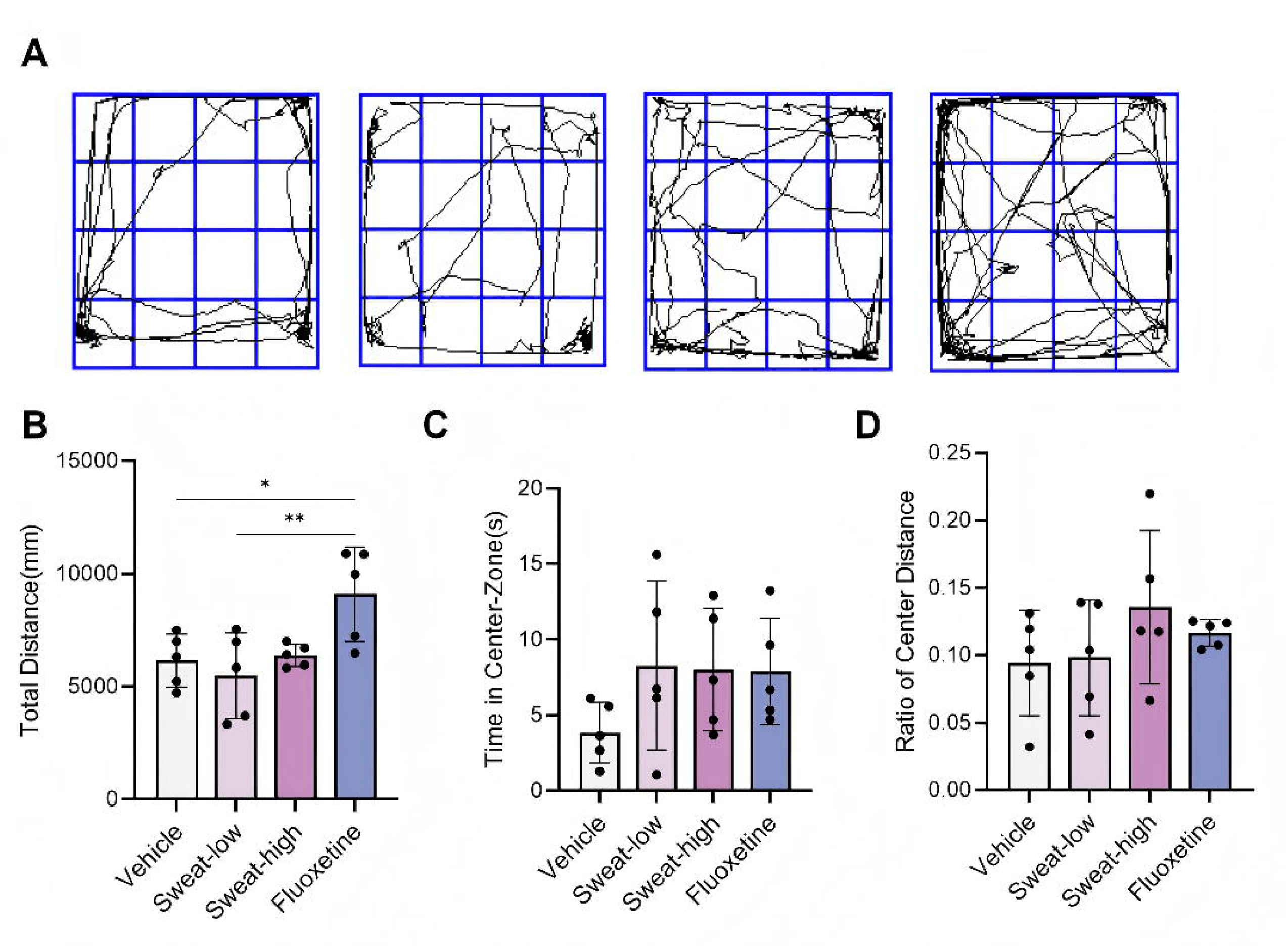
Exercise-induced sweat does not alter basal locomotor activity in the open field test (OFT). (A) Representative locomotion tracking maps of mice from each group in the OFT. Quantification of (B) total distance traveled, (C) time spent in the center region, and (D) the proportion of distance traveled in the center region relative to the total distance. Data are presented as mean ± SD (n = 5 mice/group). \**P* < 0.05, \*\**P* < 0.01.

## Discussion

Although some therapeutic agents are available for depression, clinical management still lacks safe, cost-effective, and highly efficient interventions. Therefore, identifying alternative therapies that effectively mitigate neuroinflammation and alleviate depressive behaviors is crucial to improving patient outcomes and reducing the global healthcare burden. This study represents the first to demonstrate that, in an LPS-induced depression mouse model, exercise-induced sweat treatment significantly enhances sucrose preference, reduces immobility in the TST, and lessens anxiety-like behavior in the EPM compared with the vehicle group. Notably, the therapeutic effect of high-dose sweat intervention was comparable to or even better than that of the antidepressant fluoxetine. These findings suggest that exercise-induced sweat is not merely a metabolic byproduct but also serves as a potent agent capable of mitigating depression-related neuroinflammation. This work provides a novel theoretical framework for developing sweat as a natural preparation for neuropsychiatric disorders.

Several limitations of exist in the current study and should be improved in the future. First, sweat treatment was evaluated only in a murine depression model, with analyses focused primarily on behavioral and phenotypic outcomes rather than the specific bioactive factors and molecular mechanisms involved. Future research will focus on deciphering these underlying mechanisms. Second, the sweat used in this study was collected exclusively from healthy female volunteers with a relatively small sample size. Given that sweat composition is influenced by sex, age, and individual physiological status, the generalizability of our findings may be limited^17,37,38^. Future studies should expand the donor pool to include males and different age groups to validate the consistency of the treatment’s efficacy. Additionally, this study only explored sweat induced by moderate-intensity aerobic exercise; the effects of sweat from other exercise intensities or types remain unclear. Finally, while this study established that exercise-induced sweat mitigates depressive-like behavior, its translational potential for human patients remains to be confirmed. Well-designed clinical trials are needed to evaluate the safety, efficacy, and optimal administration route of sweat in patients with depression, while accounting for comorbidities and individual differences.

## Conclusions

In summary, this study provides preliminary evidence that exercise-induced sweat treatment significantly alleviates depressive and anxiety-like behaviors in mice. Given its natural origin, inherent bioactivity, low cost, and high biocompatibility, sweat emerges as a promising candidate for developing novel, safe, and cost-effective antidepressants. Addressing current limitations through in-depth mechanistic studies, expanded donor cohorts, and translational research will be critical to advancing this innovative strategy toward clinical application, offering a new therapeutic option for the management of depression.

## Acknowledgements

This study was supported by the Natural Science Foundation of Hubei Province (2025AFD622) and the Natural Science Foundation of China [82427801, 32301239, 62025102].

## Competing interests

The authors declare no competing interests.

## Author Contributions

QC proposed the project. HY and ML determined the exercise protocol. HL designed the animal experiments. SX performed the animal experiments. HY, ML, and QF performed experiments of RNA-sequencing and reverse transcriptomics. JC, TL and BY took part in data analysis. HY, SX, ML and QC wrote the draft manuscript. CC, HL and QC supervised the study.

## Notes

https://www.ncbi.nlm.nih.gov/geo/query/acc.cgi?acc=GSE315706&token=yncbaikghturpuv

### Competing Interest Statement

The authors have declared no competing interest.

